# Recognition of cellular RNAs by the S9.6 antibody creates pervasive artefacts when imaging RNA:DNA hybrids

**DOI:** 10.1101/2020.01.11.902981

**Authors:** John A. Smolka, Lionel A. Sanz, Stella R. Hartono, Frédéric Chédin

## Abstract

The contribution of RNA:DNA hybrid metabolism to cellular processes and disease states has become a prominent topic of study. The S9.6 antibody recognizes RNA:DNA hybrids with a subnanomolar affinity, making it a broadly used tool to detect and study RNA:DNA hybrids. However, S9.6 also binds double-stranded RNA *in vitro* with significant affinity. Though frequently used in immunofluorescence microscopy, the possible reactivity of S9.6 with non-RNA:DNA hybrid substrates *in situ*, particularly RNA, has not been comprehensively addressed. Furthermore, S9.6 immunofluorescence microscopy has been methodologically variable and generated discordant imaging datasets. In this study, we find that the majority of the S9.6 immunofluorescence signal observed in fixed human cells arises from RNA, not RNA:DNA hybrids. S9.6 staining was quantitatively unchanged by pre-treatment with the human RNA:DNA hybrid-specific nuclease, RNase H1, despite experimental verification *in situ* that S9.6 could recognize RNA:DNA hybrids and that RNase H1 was active. S9.6 staining was, however, significantly sensitive to pre-treatments with RNase T1, and in some cases RNase III, two ribonucleases that specifically degrade single-stranded and double-stranded RNA, respectively. In contrast, genome-wide maps obtained by high-throughput DNA sequencing after S9.6-mediated DNA:RNA Immunoprecipitation (DRIP) are RNase H1-sensitive and RNase T1- and RNase III-insensitive. Altogether, these data demonstrate that the S9.6 antibody, though capable of recognizing RNA:DNA hybrids *in situ* and *in vitro*, suffers from a lack of specificity that precludes reliable imaging of RNA:DNA hybrids and renders associated imaging data inconclusive in the absence of controls for its promiscuous recognition of cellular RNAs.

## INTRODUCTION

RNA:DNA hybrids have emerged as a biologically interesting and potentially disease-relevant species of nucleic acid. In particular, R-loops, RNA:DNA hybrids that form via hybridization of single-stranded RNA (ssRNA) to a complementary strand of a DNA duplex, displacing the other DNA strand into a single-stranded state, have been proposed to cause DNA damage and regulate various cellular processes [1-3]. R-loop formation is thought to be a primarily co-transcriptional phenomenon [4], and elevated R-loop levels have been invoked by many studies as a link between transcription and genomic instability [5-10]. Much of the supporting evidence has relied on the use of the S9.6 mouse monoclonal antibody. S9.6 was initially reported to specifically recognize RNA:DNA hybrids [11], and has thus been widely used to isolate, sequence, measure, and image RNA:DNA hybrids in a variety of cell types from a variety of organisms [12-18].

Since the initial report on S9.6 from Bogluslawski et al. (1986), subsequent studies have shown that S9.6 can also bind double-stranded RNA (dsRNA) [19, 20] with a nanomolar affinity similar to its affinity for RNA:DNA hybrids [21]. This has made the use of ribonuclease H (RNase H) enzymes, nucleases that specifically degrade the RNA strand of RNA:DNA hybrids, critical in verifying the RNA:DNA hybrid-dependence of measurements made using S9.6-based assays [22]. While RNase H pre-treatments are routinely used as negative controls in molecular S9.6-based methods like immunoprecipitations and dot blots [23], cellular imaging using S9.6 is frequently performed with no enzymatic controls [8, 24-28]. When RNase H controls are implemented, results vary from study to study, with some studies reporting removal of S9.6 immunofluorescence signal by exogenous RNase H treatment and others finding RNase H-resistant signal [29-34]. Additionally, the S9.6 staining pattern itself varies from study to study, often coincident with methodological differences in fixation, permeabilization, and buffers used to prepare and/or enzymatically treat cells prior to immunolabeling [8, 24, 29, 30, 32, 35, 36]. Lastly, though a common goal of using S9.6 is to image R-loop structures in the nucleus, cytoplasmic S9.6 signal has been consistently observed across studies (see references above). This signal is often unaddressed or attributed to R-loops arising from the mitochondrial genome [13, 28]. However, conclusive experimental evidence to establish the origin of this signal and its sensitivity to exogenous RNase H treatment is lacking.

In this study, we established a protocol to test the RNA:DNA hybrid- and RNA-dependence of S9.6 staining using structure-specific nucleases to selectively and separably degrade RNA:DNA hybrids and RNA. In addition, we implemented a positive control for S9.6 staining and *in situ* RNase H1 activity using synthetic Cy5-labeled RNA:DNA hybrids transfected into human cells prior to imaging. Using this approach, we verified that exogenous RNA:DNA hybrids can be recognized by S9.6 and degraded by RNase H1 *in situ* under methanol-fixed conditions. However, we demonstrate that the endogenous structures labeled by S9.6 in normally cultured cells are resistant to pre-treatments with RNase H1. In contrast, S9.6 labeling was significantly reduced by pre-treatments with enzymes that specifically degrade ssRNA and, to a lesser extent, dsRNA. Thus, we conclude that images obtained through immunofluorescence microscopy using the S9.6 antibody are vulnerable to artefactual signal that reflects cellular RNA content, not RNA:DNA hybrids.

## MATERIALS AND METHODS

### Cell culture

U2OS and HeLa cells were obtained from ATCC and grown in high-glucose Dulbecco’s Modified Eagle’s medium (DMEM) supplemented with 10% fetal bovine serum (FBS) and 1% penicillin / streptomycin. Samples were seeded with cells at equal densities 1-2 days prior to experiments and were grown to 40-60% confluency. Cells were regularly tested and verified to be negative for mycoplasma prior to experiments.

### Fixation and labeling for immunofluorescence

All cell culture samples were grown, fixed, permeabilized, washed, enzymatically treated, immunostained, and imaged in 35 mm glass bottom poly-D-lysine-coated dishes (P35GC-1.5-14-C, MatTek) using 2 mL volumes of media and buffer solutions, unless indicated otherwise. All steps for fixation and immunofluorescence were carried out at room temperature, unless indicated otherwise. For methanol fixation, cells were fixed with 2 mL of ice-cold, 100% methanol for 10 minutes at −20°C. Cells were then washed once with PBS before subsequent preparation steps for immunofluorescence. Permeabilization steps were unnecessary and omitted for methanol-fixed samples. For formaldehyde fixation, cells were fixed in freshly prepared 1% formaldehyde in PBS for 10 minutes. Fixation solutions were quenched with the addition of 200 μL of PBS with 1 M glycine. Formaldehyde-fixed samples were washed once with PBS, and then incubated in permeabilization buffer (PBS with 0.1% Triton X-100) for 10 minutes. For both methanol- and formaldehyde-fixed samples, samples were then incubated in staining buffer (TBST with 0.1% BSA (A9647-50G, Sigma)) for 10 minutes with rocking. Enzymatic treatments were done in staining buffer supplemented with 3 mM magnesium chloride with 1:200 dilutions of RNase T1 (EN0541, Thermo Fisher), RNase III (ShortCut RNase III, M0245S, New England Biolabs), and/or human RNase H1 [37] and incubated with rocking for 1 hour. Samples were subsequently washed by incubating with staining buffer for 10 minutes with rocking. For primary immunolabeling, samples were incubated in staining buffer with a 1:1000 dilution of rabbit anti-HSP27 (06-478, Millipore) for cell body labeling and/or mouse S9.6 (purified from the HB-8730 hybridoma cell line) for a minimum of 1 hour at room temperature or overnight at 4°C with rocking. Samples were then washed once with staining buffer and incubated with 1:2000 dilutions of secondary anti-mouse AlexaFluor 488 conjugate (A28175, Invitrogen) and/or anti-rabbit AlexaFluor 594 conjugate (A11037, Life Technologies) in the same manner as for the primary antibody incubation, with samples kept concealed from light from this step onward. Samples were then incubated with a 2.5 μg/mL DAPI dilution in staining buffer for 1-2 minutes and washed in TBST for 10 minutes with rocking before being stored in PBS at 4°C until imaging. Samples were imaged in PBS. To label mitochondria, live cells were stained with 500 nM MitoTracker Deep Red FM (Thermo Fisher) in complete medium for 15 min at 37°C, and then incubated in MitoTracker-free complete medium for 15 minutes prior to fixation. For each experiment, all samples were prepared, treated and stained in parallel from one master enzyme, antibody, and/or dye dilution to ensure uniform treatment and staining efficiencies.

### Oligonucleotide preparation and transfection

Oligos were synthesized and HPLC-purified by Integrated DNA Technologies. Lyophilized DNA and RNA oligos were resuspended in autoclaved nanopure water to make 100 μM stocks. To prepare substrates for enzymatic tests and transfections, DNA and/or RNA oligos were annealed in TBST at 10 μM concentrations by heating to 95°C for 3 minutes and then cooling to room temperature. Transfections were performed using Lipofectamine 2000 (Thermo Fisher) at a 1:50 dilution in 200 μL of Opti-MEM (Gibco) with a 1:100 dilution of RNA:DNA hybrids. The lipofectamine/RNA:DNA hybrid mix was then added dropwise to cells in 2 mL of complete media. Transfections were carried out for 3 hours and cells were washed once with warm complete medium and incubated for an additional 6 hours before fixation.

The following oligonucleotide sequence, along with its complement and cognate RNA sequences, was used to design all nucleic substrates used: 5’-AGCTATAGTGACTGACGTTATCATGATGCTAGAGTCTCGATCGATAGTGTAGCT-3’

### Imaging and image analysis

Fixed cell imaging was performed using the spinning-disk module of an inverted objective fluorescence microscope [Marianas spinning disk confocal (SDC) real-time 3D Confocal-TIRF (total internal reflection) microscope; Intelligent Imaging Innovations] with a 63X or 100X objective. For each experiment, all conditions were imaged in parallel with identical exposure times and laser settings. Images were analyzed and quantified using ImageJ/FIJI programs, and statistics and data visualization were done using RStudio. P-values were determined by a Wilcoxon Mann-Whitney test with the R wilcox.test() function. Whole cell and nuclear mean S9.6 intensities were determined from a single plane for individual images. Whole cell and nuclear regions were defined by thresholding or manual tracing using HSP27 and DAPI, respectively. For quantification of Cy5-associated S9.6 mean intensities, the Cy5 channel of each analyzed single plane image was manually thresholded to generate a binary image. Analyze Particles was applied to convert binary Cy5 images into Regions of Interest (ROIs) used to quantify the S9.6 intensities of Cy5-occupied regions of an image. Each ROI was defined as an individual Cy5 particle.

### Enzyme validation and gel electrophoresis

Single- and double-stranded oligonucleotide substrates were diluted in TBST with 0.1% BSA and 3 mM magnesium chloride to 1 μM final concentrations Previously mentioned enzymes and RNase A (EN0531, Thermo Fisher) were used at a 1:200 final dilution and incubated with indicated nucleic acid substrates for 1 hour at room temperature. To test ShortCut RNase III under manganese-supplemented conditions, reactions were carried out in 50 mM Tris-HCl pH 7.5, 75 mM KCl and 3 mM MnCl_2_ with 0.1% BSA. After incubation, a 1/10th volume of 50% glycerol was added to each sample before gel electrophoresis through 10% polyacrylamide TBE gels run at 80V for 45 minutes. Gels were post-stained with ethidium bromide and washed in TBE before imaging.

### DNA:RNA Immunoprecipitation (DRIP) followed by DNA sequencing

DNA:RNA immunoprecipitation was performed as previously described [38] with modifications. Briefly, nucleic acids gently extracted from unfixed NTERA-2 cells were sheared by sonication (12 cycles at high power, 15s ON / 90s OFF) with a Bioruptor (Diagenode). 4.4 μg of sheared nucleic acids were incubated with 10 μg of S9.6 antibody for immunoprecipitation overnight at 4°C. To build strand-specific sequencing libraries, immunoprecipitated nucleic acids were subjected to a second-strand DNA synthesis step in which dUTP was used instead of dTTP. After verifying the quality of immunoprecipitation using qPCR at standard positive and negative loci [38], strand-specific libraries were constructed, and DNA was sequenced using the HiSeq4000 platform. For RNase treatments, 15 μg of sheared DNA were treated with RNase H1, RNase T1 and/or RNase III for 1 hour at 37°C using a 1/100 dilution of the stock enzymes in 50 mM Tris-HCl pH 7.5, 75 mM KCl and 3 mM MgCl_2_. DNA was then purified with phenol/chloroform and ethanol precipitated before being used in S9.6 immunoprecipitations.

### Normalization, peak calling and data analysis for DRIP

Reads were mapped to the human genome using Bowtie2 [39]; only concordantly mapped reads were considered in case of paired-end samples. Normalization was performed based on total uniquely mapped reads with background correction. Macs was used to call peaks with default parameters except for –nomodel [40]. For comparison of DRIP signals across treatments, we counted read distributions across 10 kb genomic bins through the human genome. Pearson correlations were then systematically performed between each sample, excluding self-correlations. P-values were calculated using the Wilcoxon Mann-Whitney test with the R wilcox.test() function. For XY plots, the averaged DRIP signal was measured over 10 kb bins for each treatment group (RNase T1, RNase III, RNase T1+III, and RNase H; y-axis) and compared to averaged DRIP signal observed in the mock treated samples (x-axis). Their linear regression values were then calculated using the R lm() function.

## RESULTS

### RNA constitutes the majority of the S9.6 immunofluorescence signal in the cytoplasm and nucleus of human cells

Using human U2OS cells fixed with methanol, the most commonly used fixative for S9.6 staining, S9.6 immunofluorescence (IF) signal was observed in both the nucleus and the cytoplasm (Figure 1A). Nuclear staining was predominantly nucleolar in morphology and localization. These staining patterns are congruent with several previous reports [26, 28, 30, 41] and have been explained by claims that ribosomal DNA is a hotspot for R-loop formation in the nucleus [42] and that mitochondrial genomes in the cytoplasm contain R-loops [43]. To determine if the cytoplasmic S9.6 signal derives from mitochondria, we labeled mitochondria prior to fixation using the vital dye, MitoTracker Deep Red FM, and assessed the degree of co-localization with S9.6. Cytoplasmic S9.6 staining had a distribution more similar to HSP27, which labels the entire cell body, than to mitochondria (Figure 1B), and S9.6 immunofluorescence signal did not show enrichment in or correlation with mitochondrial territories (Figure 1C). These observations agree with prior work in human cells grown under standard conditions [32], and indicate that mitochondria are not the source of the majority of cytoplasmic S9.6 staining, and that S9.6 detects extra-mitochondrial structures throughout the cytoplasm.

**FIGURE 1:**
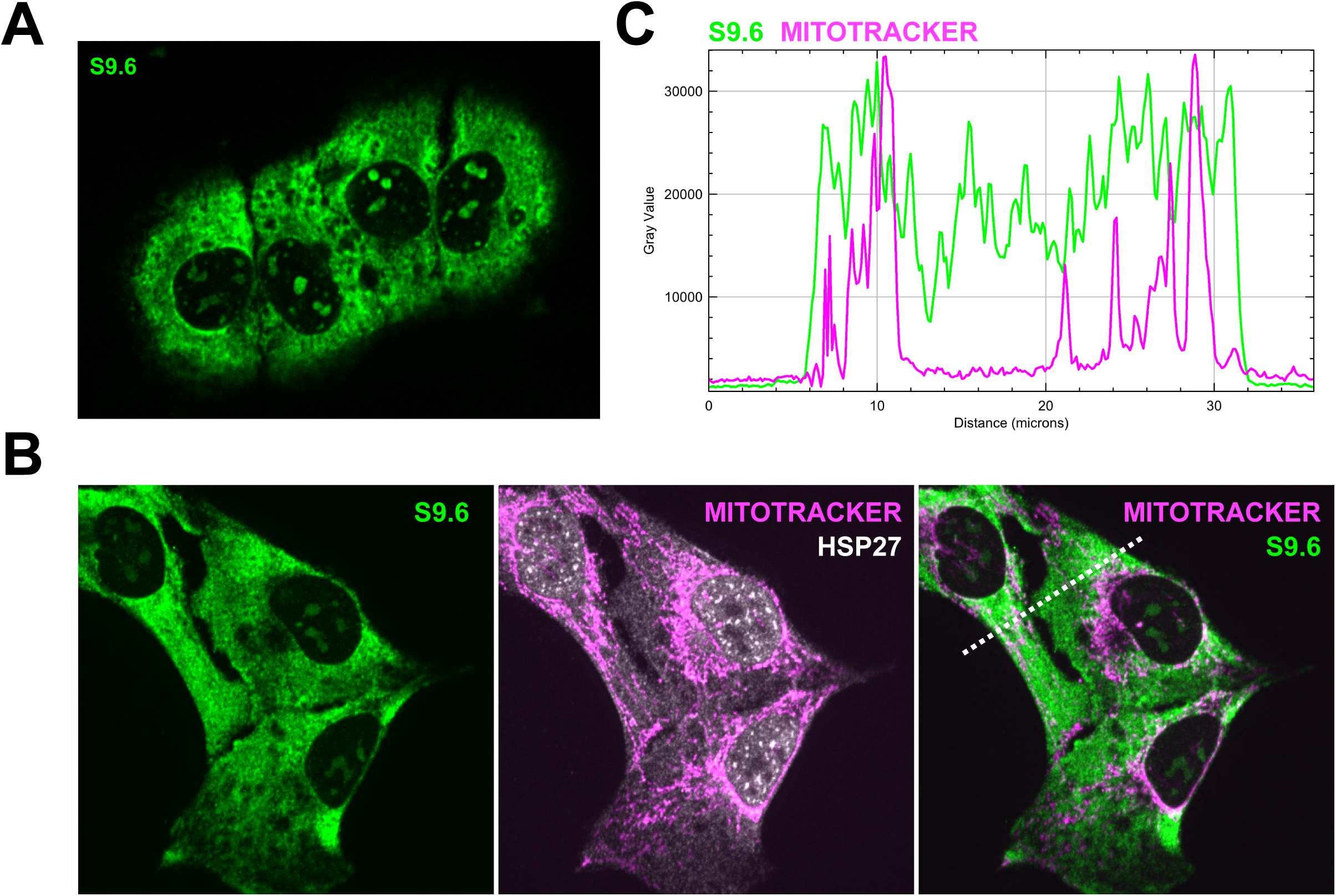
**A.** A single plane confocal immunofluorescence image of methanol-fixed U2OS cells stained with S9.6 (green). **B.** Maximum intensity projection (MIP) of U2OS cells labeled with MitoTracker Deep Red FM (magenta) and stained with S9.6 (green) and anti-HSP27 (white). **C.** Line scan plotting the intensity values for S9.6 and MitoTracker Deep Red FM in cells from the MIP in B.

Informed by previous reports showing S9.6 can recognize dsRNA *in vitro*, we hypothesized that various species of cellular RNA in fixed cells may be the source of cytoplasmic S9.6 signal, and possibly nuclear signal as well. To test this hypothesis, we identified ribonucleases that could robustly degrade RNA without degrading RNA:DNA hybrids. Under a magnesium-supplemented condition, which is necessary for RNase H activity [44], we verified that commercial RNase T1 and RNase III enzymes were capable of specifically and efficiently degrading ssRNA and dsRNA substrates. Importantly, under the same condition, these enzymes had negligible activity on RNA:DNA hybrid substrates (Figure S1A) and full-length, recombinantly-purified human RNase H1 [37] efficiently digested the RNA strand of the RNA:DNA hybrid substrate (Figure S1B). However, RNase A degraded RNA:DNA hybrids within 1 hour at room temperature (Figure S1C), indicating it was not a suitable enzyme to use for specific degradation of RNA. Additionally, RNase III degraded RNA:DNA hybrids in the presence of manganese (Figure S1D), indicating that the manganese supplementation recommended by New England Biolabs could not be used.

We tested the effects of validated enzymes *in situ* on S9.6 staining by subjecting fixed samples to mock or enzymatic treatments prior to immunolabeling under the same conditions used for the *in vitro* tests above. As expected, mock-treated methanol-fixed cells stained with S9.6 showed pan-cytoplasmic signal and a nucleolar staining pattern within the nucleus (Figure 2A). The average nuclear S9.6 intensity was 35.4% of the average total cellular intensity (Figure 2B), indicating that cytoplasmic signal was the predominant constituent of the total cellular S9.6 signal. Pre-treatment with RNase H1 did not have a significant effect on S9.6 staining in the cytoplasm or the nucleus (Figure 2, A and B). Pre-treatment with RNase III had minimal effect on total cellular signal (9.6% reduction, p-value: 0.17) concomitant with a minor but significant effect on nuclear signal (20.1% reduction, p-value: 0.002) (Figure 2B). In comparison, pre-treatment with RNase T1 led to a significantly larger decrease in both nuclear and cytoplasmic S9.6 staining (Figure 2A), reducing total cellular S9.6 staining by 58.9% (p-value: 9.4e-12) (Figure 2B). Notably, RNase T1 treatment reduced average nuclear IF signal by 62.7% (p-value: 8.9e-15) and led to a loss of nucleolar staining (Figure 2, A and B). These effects were consistent across biological replicates and also observed in HeLa cells (Figure S2, A and B). Since it was previously reported that S9.6 labeling of cells may be limited by formaldehyde fixation [24], we also imaged formaldehyde-fixed U2OS cells. Application of S9.6 IF and RNase T1 and III pre-treatments to formaldehyde-fixed cells largely recapitulated findings using methanol-fixed cells, indicating that formaldehyde fixation does not significantly alter the nature of S9.6 staining (Figure S3, A and B).

**FIGURE 2:**
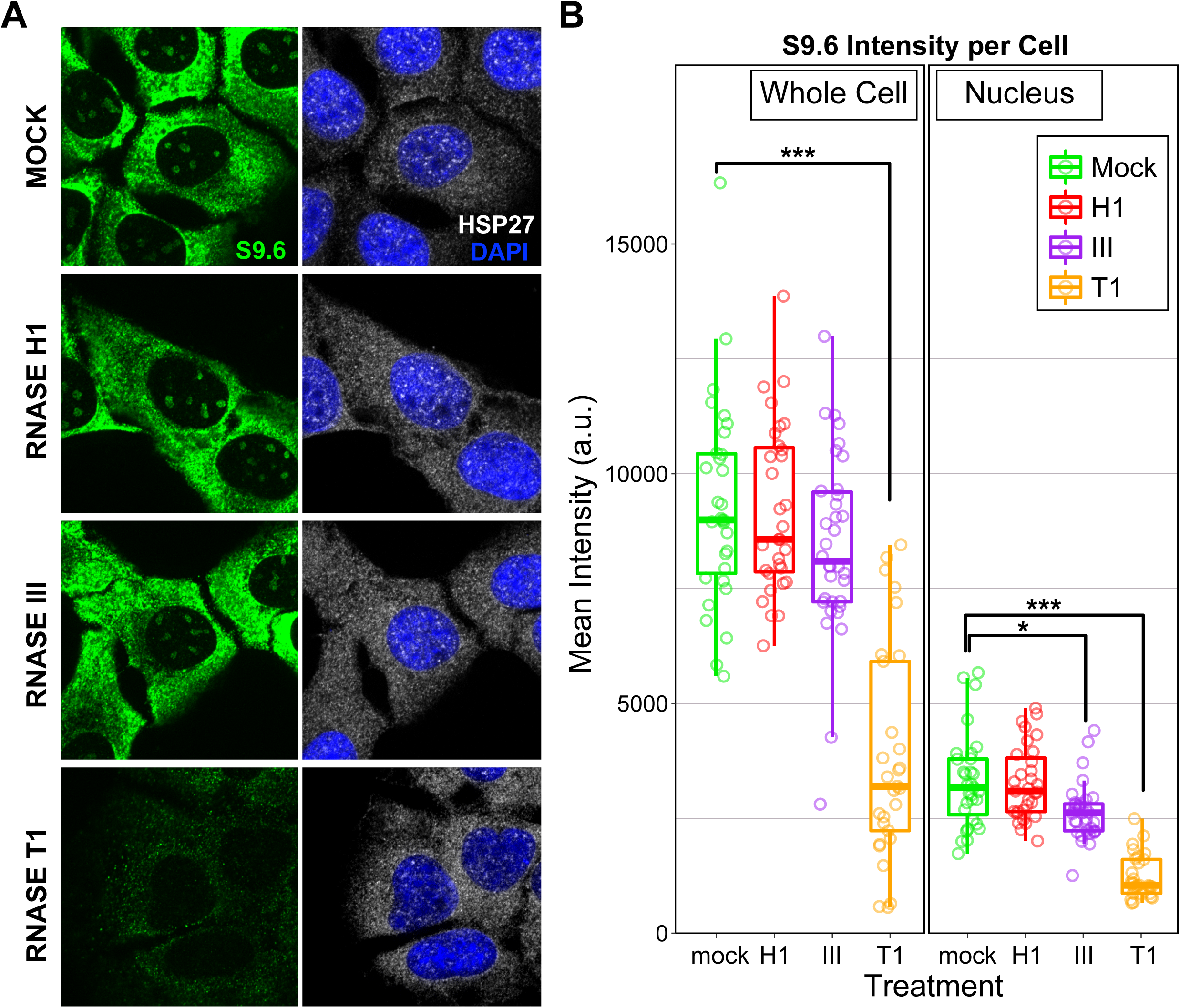
**A.** Representative images of single planes of cells that were mock-treated or pre-treated with RNase H1, RNase III, and RNase T1 for 1 hour at room temperature. **B.** Quantification of whole cell and nuclear mean S9.6 intensities for individual cells that were mock- or enzyme-treated. Plots represent combined data from two biological replicates.

After treatment with a combination of RNase T1 and RNase III, residual RNase H1-insensitive cytoplasmic S9.6 signal was observed (Figure S4). We hypothesize that this signal corresponds to RNase T1- and RNase III-resistant RNAs. It is important to note that RNase T1 is an endonuclease that specifically cleaves ssRNA at guanine residues, and that RNase III efficiently fragments dsRNA rather than hydrolyzing it completely under these conditions (Figure S1A). Thus, these enzymes may have inherent limitations in their abilities to fully degrade all RNA species due to their unique enzymatic behaviors. Regardless, these observations taken in total support that the majority of S9.6 signal in methanol-fixed cells, both in the nucleus and the cytoplasm, stems from the binding of S9.6 to RNA, not RNA:DNA hybrids.

### The S9.6 antibody can detect transfected RNA:DNA hybrids that are degraded by treatment with exogenous RNase H1 *in situ*

Given the negative results obtained above when testing for the RNA:DNA hybrid-dependence of S9.6 IF signal, we sought a positive control for detection of RNA:DNA hybrids by S9.6 and degradation of RNA:DNA hybrids by RNase H1 *in situ*. Using 54 base-pair RNA:DNA hybrids that were fluorescently-labeled with a 5’-Cy5 modification on the DNA strand, we transfected U2OS cells prior to fixation and again subjected samples to mock or RNase H1 treatments after fixation and prior to immunolabeling (Figure 3A). After lipofection with Cy5-labeled RNA:DNA hybrids, Cy5 foci were observed in and/or on the cell body of fixed cells (Figure 3B). Consistent with prior results [45], Cy5 foci regularly appeared to be in endosomal-like compartments that excluded HSP27 (Figure 3B, inset), suggesting transfected hybrids had been internalized. Staining with S9.6 revealed prominent S9.6 foci that overlapped Cy5 foci (Figure 3, B and C). As expected, there was also pan-cytoplasmic and nucleolar S9.6 IF signal that was independent of the focal Cy5 signal (Figure 3, B and C). Transfection with unhybridized, single-stranded Cy5-labeled DNA oligos produced Cy5 foci with no accompanying S9.6 foci (Figure S5; we note that ssDNA oligos displayed more efficient nuclear accumulation than RNA:DNA hybrids), further supporting that the S9.6 foci arising from transfection with hybridized oligos were RNA:DNA hybrid-dependent. These observations support that exogenous hybrids can be used *in situ* as a positive control for S9.6 immunolabeling.

**FIGURE 3:**
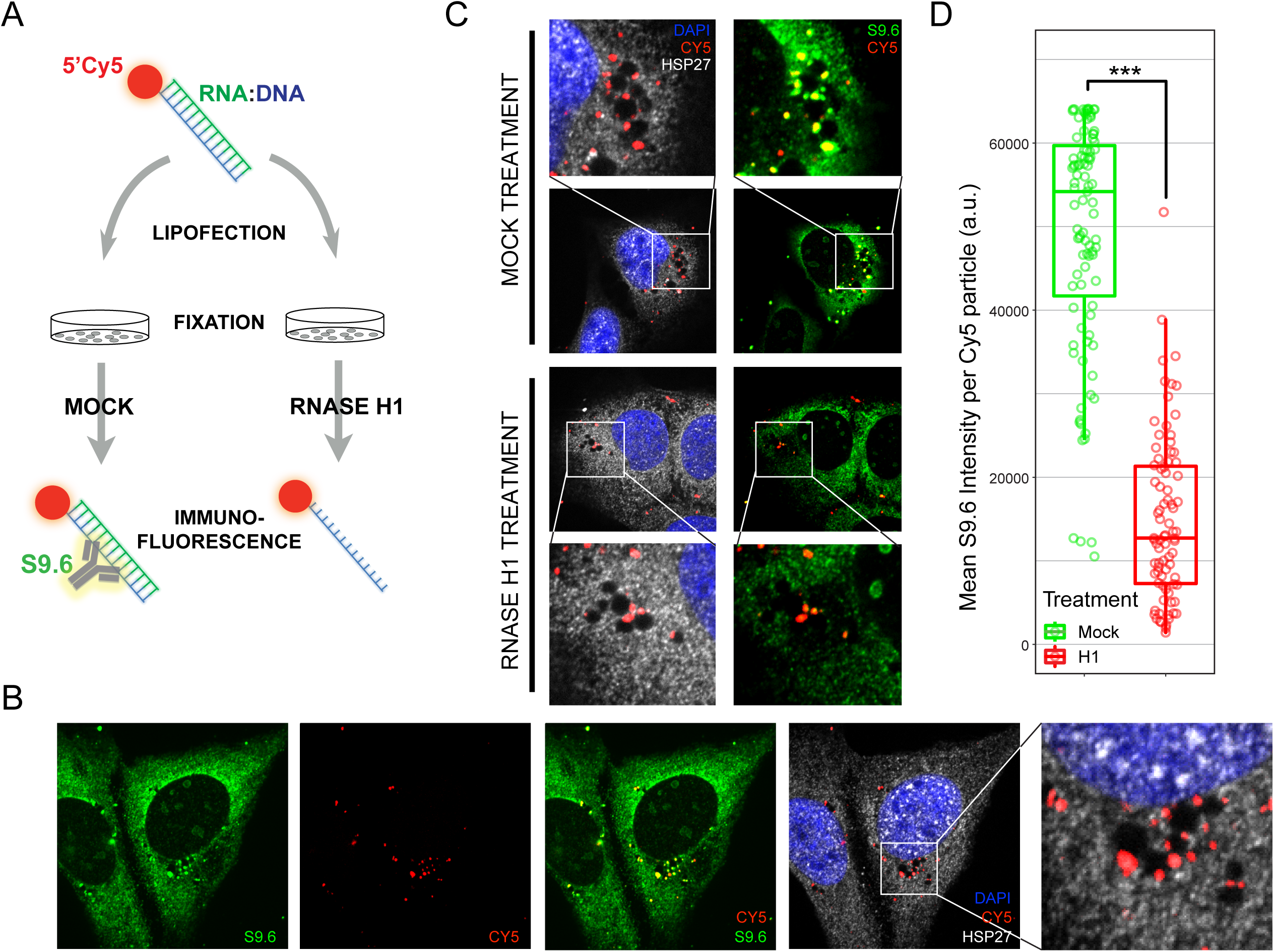
**A.** Schematic of the RNA:DNA hybrid transfection strategy used as a positive control for *in situ* S9.6 staining and RNase H1 enzymatic activity. **B.** Representative single plane image showing transfected U2OS cells containing Cy5-labeled RNA:DNA hybrids (red) that co-localize with S9.6 (green). **C.** Representative single plane images of mock- and RNase H1-treated transfected cells. **D.** Quantification of the mean S9.6 intensities of individual Cy5 foci in mock- and RNase H1-treated cells. Plots represent combined data from two biological replicates.

To verify that exogenous RNase H1 treatment could effectively degrade hybrids *in situ* in a methanol-fixed environment, we compared the overlapping S9.6 intensities of individual Cy5 foci in transfected cells that were mock- and RNase H1-treated (Figure 3, C and D). RNase H1 treatment reduced the average Cy5 focus-associated S9.6 signal by 69.7% (p-value: 4e-26) (Figure 3D), while the Cy5-independent cytoplasmic and nuclear signal persisted. This observation demonstrates that RNase H1 is active under these conditions and capable of efficiently degrading RNA:DNA hybrids *in situ*. Additionally, Cy5-associated S9.6 foci persisted after treatment with a combination of RNases T1 and III, while the Cy5-independent cytoplasmic and nuclear signal was reduced (Figure S5), further supporting that RNases T1 and III selectively degrade RNA that is detected by S9.6. Altogether, these results confirm S9.6 can detect *bona fide* RNA:DNA hybrids in fixed cellular environments and that RNase H1 can degrade these substrates post-fixation. However, this indicates that S9.6 IF signal observed using normally cultured human cells is resistant to RNase H1 treatments because S9.6 is predominantly labeling RNA, and not because of insufficiencies in RNase H1 treatment.

### S9.6 immunoprecipitation followed by high-throughput DNA sequencing is insensitive to RNase T1 and RNase III pre-treatments

Since S9.6 has been widely used as a genomics tool to map R-loops, we tested the effects of RNase T1, RNase III, and RNase H1 pre-treatments on DNA:RNA hybrid ImmunoPrecipitation (DRIP) using S9.6 followed by high-throughput, strand-specific DNA sequencing. For systematic comparison, sonicated, unfixed nucleic acids extracted from human NTERA-2 cells were mock- or enzyme-treated, using the same enzymes used in the controls for IF with similar buffer conditions, prior to immunoprecipitation with S9.6. Genome-wide maps obtained via sonication DRIP-seq (sDRIP-seq) showed predominantly genic, strand-specific signals associated with the direction of transcription (Figure 4A). This signal was abrogated by RNase H1 treatment, supporting that DRIP recovers transcription-dependent RNA:DNA hybrids. Genome-wide analysis of sDRIP-seq maps showed that samples pre-treated with RNase T1, RNase III, or a combination of the two enzymes had consistent average correlations of 0.8 or higher with the mock-treated control (Figure 4B). By contrast, RNase H1-treatment significantly reduced the average correlation with the mock-treated condition (p-value: 0.0009). This was further reflected in aggregate analyses of sDRIP signal over transcription start sites, gene bodies, and transcription termination sites of human genes, which had stereotypical enrichments and profiles that were not altered significantly by pre-treatments with RNase T1 and/or RNase III, but were lost with pre-treatment using RNase H1 (Figure 4C). In summary, sDRIP-seq is sensitive to RNase H1 treatment, but resistant to RNase T1 and RNase III treatments, supporting the RNA:DNA-hybrid dependence of genomic profiles generated via S9.6-based DRIP followed by high-throughput DNA sequencing.

**FIGURE 4:**
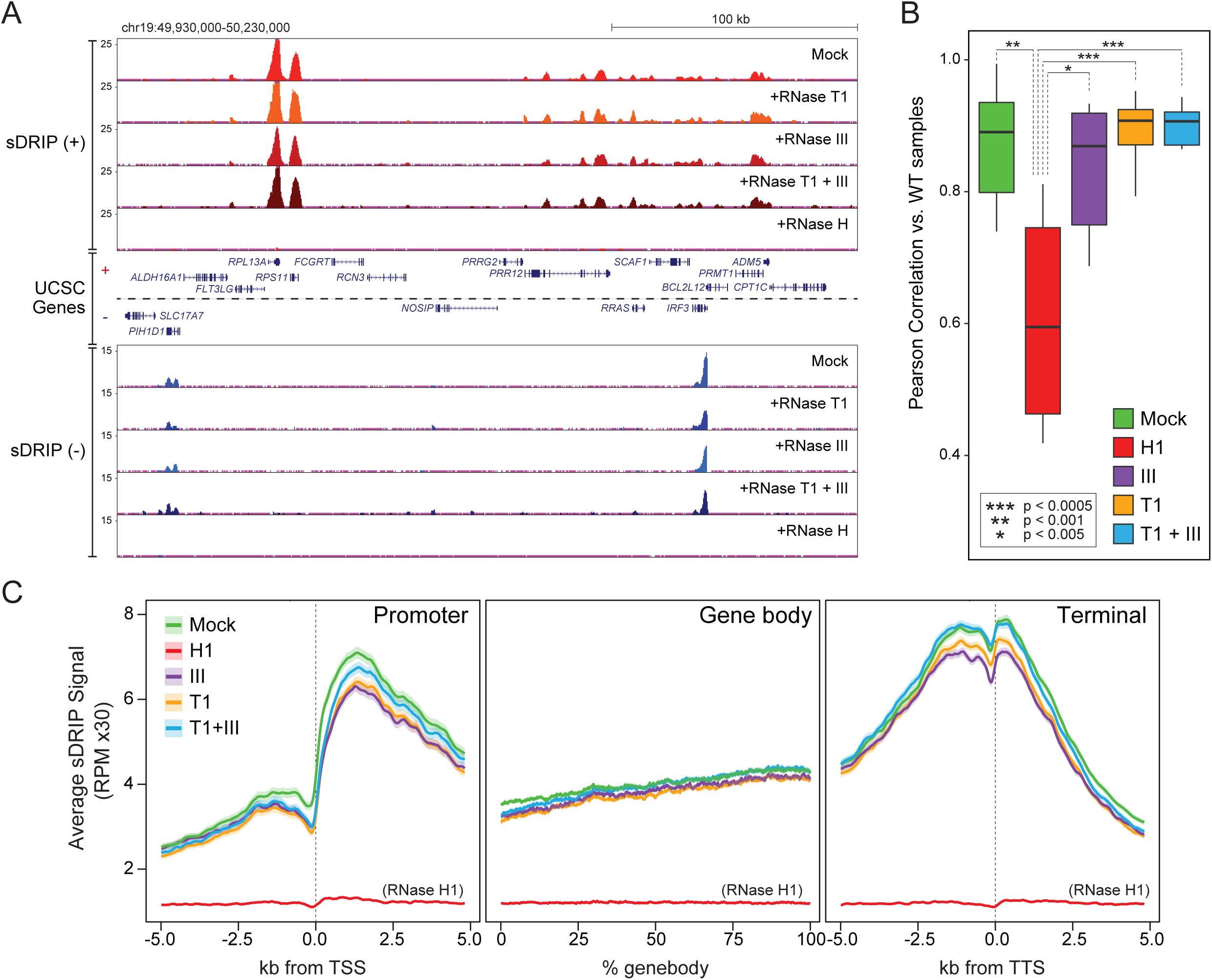
**A.** Genome browser tracks of a representative region of the human genome showing plus and minus strand sDRIP-seq signal obtained from mock-, RNase III-, RNase T1-, RNase T1 and III-, and RNase H1-treated DRIP samples. **B.** Boxplots showing the mean Pearson’s correlations of sDRIP-seq signal between mock- and enzyme-treated samples. Correlation values were calculated from data from two replicates for each condition. **C.** Metaplots of sDRIP-seq signal over the transcription start site (TSS), gene body, and transcription termination site (TTS) of genes with RNA expression levels in the top 10% of expressed genes for mock- and enzyme-treated samples. For the TSS and TTS, the signal was plotted over a +/− 5kb region. For gene bodies, the signal is shown as a percentile plot. Metaplots represent data from two replicates for each condition. Lines represent trimmed means and accompanying shaded areas represent standard error.

## DISCUSSION

We find images obtained through immunofluorescence microscopy using the S9.6 antibody predominantly reflect cellular RNA content in normally cultured human cells. This conclusion is based on two key observations: (i) S9.6 staining is sensitive to treatments with ssRNA-and dsRNA-specific nucleases; (ii) S9.6 staining is insensitive to RNase H treatment under conditions where RNase H is active *in situ*. The observation that the S9.6 antibody pervasively detects cellular RNA when imaging human cells is consistent with reports that S9.6 is not RNA:DNA hybrid-specific and has significant affinity for dsRNA. It is likely that the predominant cytoplasmic and nucleolar staining of S9.6 observed here and in other studies reflects large pools of RNAs with duplexed structural features for which S9.6 has an affinity. We note that lower cytoplasmic S9.6 staining has been observed in some studies [8, 35, 46], differing from the observations made here. These differences can be attributed to the use of various pre-extractions, chemical treatments, and washing steps prior to immunolabeling that we did not employ. Regardless of differences in sample preparation, the S9.6 antibody is liable to recognize cellular RNAs and the RNA:DNA hybrid-dependence of an observed staining pattern must be verified.

While this work used normally cultured, unperturbed human cells, many studies have used S9.6-based imaging to measure changes in RNA:DNA hybrid content under various experimental conditions. An array of gene perturbations and drug treatments have been reported to cause changes in S9.6 IF signal attributed to altered R-loop metabolism or RNA:DNA hybrid levels [8, 30, 34-36, 47-50]. Many of the genes that have been ascribed roles in R-loop regulation participate in diverse RNA metabolic processes like transcription, RNA splicing, RNA processing, and RNA export [6]. Since R-loops can form co-transcriptionally, it is conceivable that alterations in transcription and processes intimately associated with it could affect R-loop formation. However, alterations in RNA metabolism may also change the overall levels or sub-cellular distribution of various RNA species, which could in turn alter S9.6 staining. In the absence of controls for off-target RNA recognition by S9.6, such altered staining patterns could be incorrectly interpreted as a change in RNA:DNA hybrids. It may be prudent to use the controls outlined in this study to validate previous observations made using S9.6 IF.

In contrast to imaging, the sequencing of RNA:DNA hybrids to map R-loops in human cells using sDRIP-seq is not significantly affected by the off-target recognition of RNA by S9.6. When DNA, not RNA, is the substrate for sequencing, such as in DRIP-seq [13] and in sDRIP-seq (this study), the ability of RNA to contribute off-target signal is essentially eliminated. However, for DRIP-based methods that build sequencing libraries from RNA, such as in DRIPc-seq [4], caveats due to non-specific RNA-binding by S9.6 exist. In fission yeast, for example, high-abundance dsRNA species that were insensitive to RNase H pre-treatment were detected by DRIPc-seq [19]. Pre-treatment with RNase III corrected this issue, supporting that enzymatic controls that selectively degrade RNA can improve the reliability of S9.6-based genomic approaches that rely on sequencing RNA-derived materials. We suggest S9.6-based R-loop mapping should employ sequencing strategies that query R-loops through the DNA moieties of RNA:DNA hybrids rather than RNA to avoid any issue that may arise from the off-target RNA-binding of S9.6.

We note that co-immunoprecipitations using S9.6 to assay RNA:DNA hybrid-protein interactions may also be vulnerable to isolation of RNA-protein interactions. For instance, an S9.6-based proteomic screen aimed at immunopurifying RNA:DNA hybrid-associated proteins identified a large number of RNA-binding proteins. Additionally, the RNA Recognition Motif (RRM), a well-characterized ssRNA-binding motif, was the most enriched protein domain in this dataset [51]. This highlights the potential for the off-target recognition of RNA by S9.6 to be a systemic issue. To mitigate the potential for RNA to interfere with S9.6-based approaches, degradation of RNA using a standard enzyme like RNase A is an intuitive solution. However, we find the RNA:DNA hybrid-degrading activity of RNase A makes use of the enzyme a suboptimal and unreliable control. While the use of RNase III rather than RNase A is an ideal way to specifically degrade dsRNA, we find RNase III alone does not degrade a large portion of the RNAs that S9.6 can recognize in the context of imaging. Thus, to specifically and maximally degrade RNA while preserving RNA:DNA hybrids, we recommend systematic use of RNase III and RNase T1 treatments, as RNase T1 appears to specifically degrade a large portion of cellular RNAs that RNase III cannot. These pre-treatments, coupled with RNase H controls, should help investigators use S9.6 in a thoroughly controlled manner that can account for the capacity of S9.6 to recognize RNA. Ultimately, we hope that the observations made and controls established in this study can help ensure robust and accurate insights into RNA:DNA hybrid metabolism through informed use of the S9.6 antibody.

## SUPPLEMENTAL FIGURE LEGENDS

**FIGURE S1:**
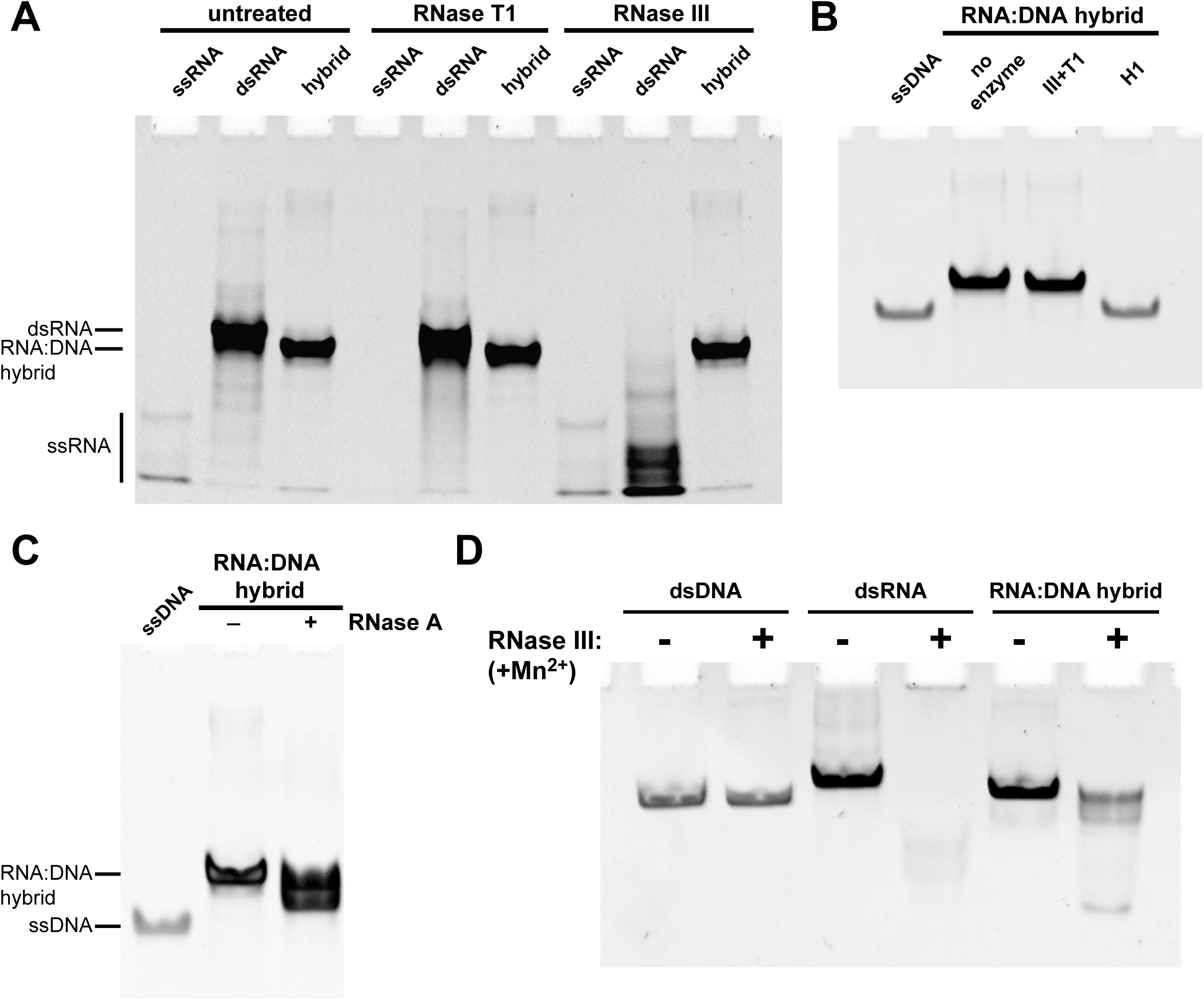
**A.** Ethidium bromide-stained polyacrylamide gels showing 54 nucleotide ssRNA and 54 basepair dsRNA and RNA:DNA hybrid substrates of the same sequence untreated and treated with RNase T1 and RNase III. Treatments were done for 1 hour at room temperature. **B.** RNA:DNA hybrids subjected to treatment with a combination of RNase T1 and III and treatment with RNase H1. **C.** Treatment of RNA:DNA hybrid substrates with RNase A at 0.05 mg/mL. **D.** Treatment of dsDNA, dsRNA, and RNA:DNA hybrids with ShortCut RNase III under manganese-supplemented conditions.

**FIGURE S2:**
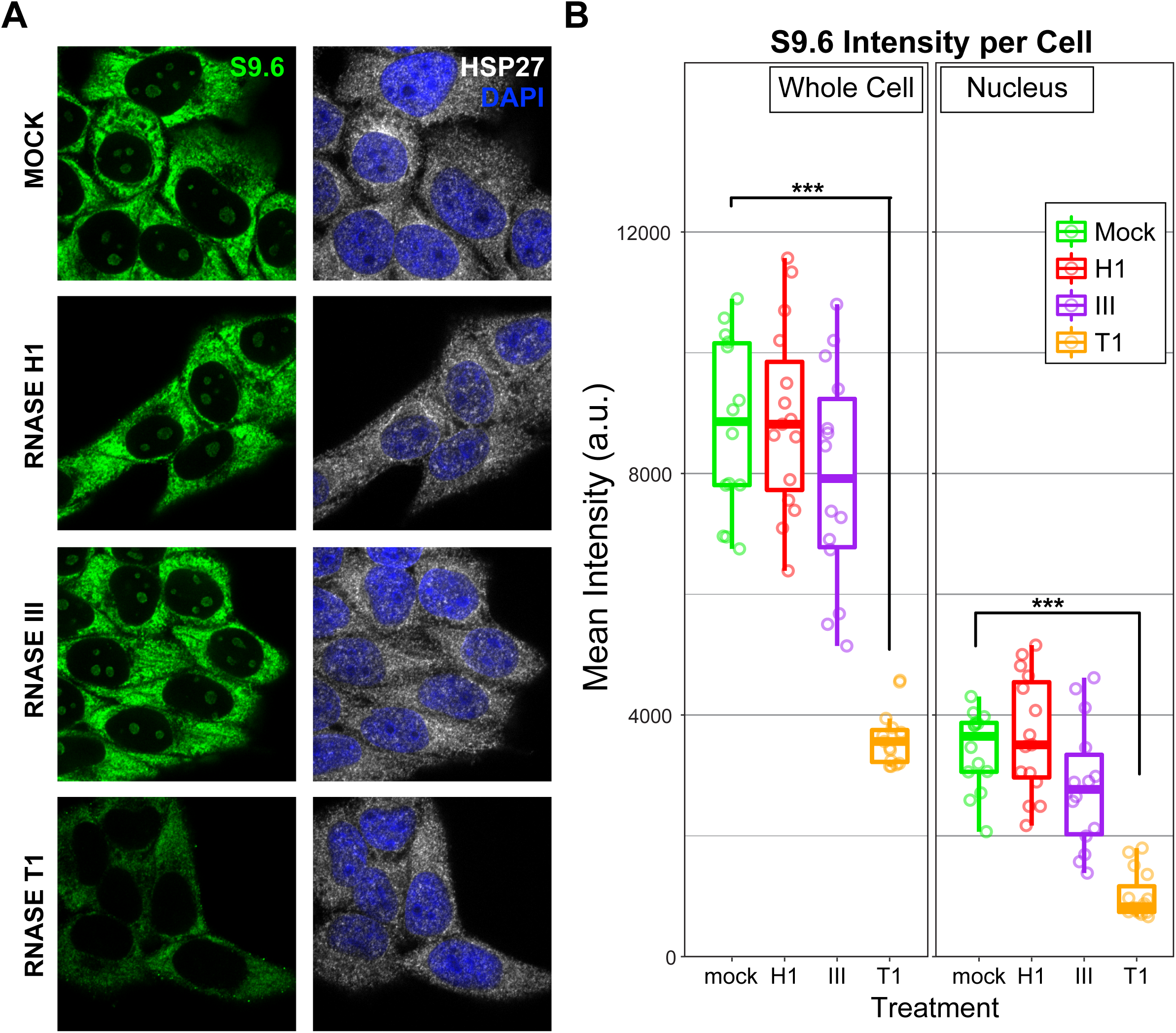
**A.** Representative images of single planes of HeLa cells that were mock-treated or pre-treated with RNase H1, RNase III, and RNase T1 post-fixation for 1 hour at room temperature and stained with S9.6 (green) and anti-HSP27 (white). **B.** Quantification of whole cell and nuclear mean S9.6 intensities for individual cells that were mock- or enzyme-treated.

**FIGURE S3:**
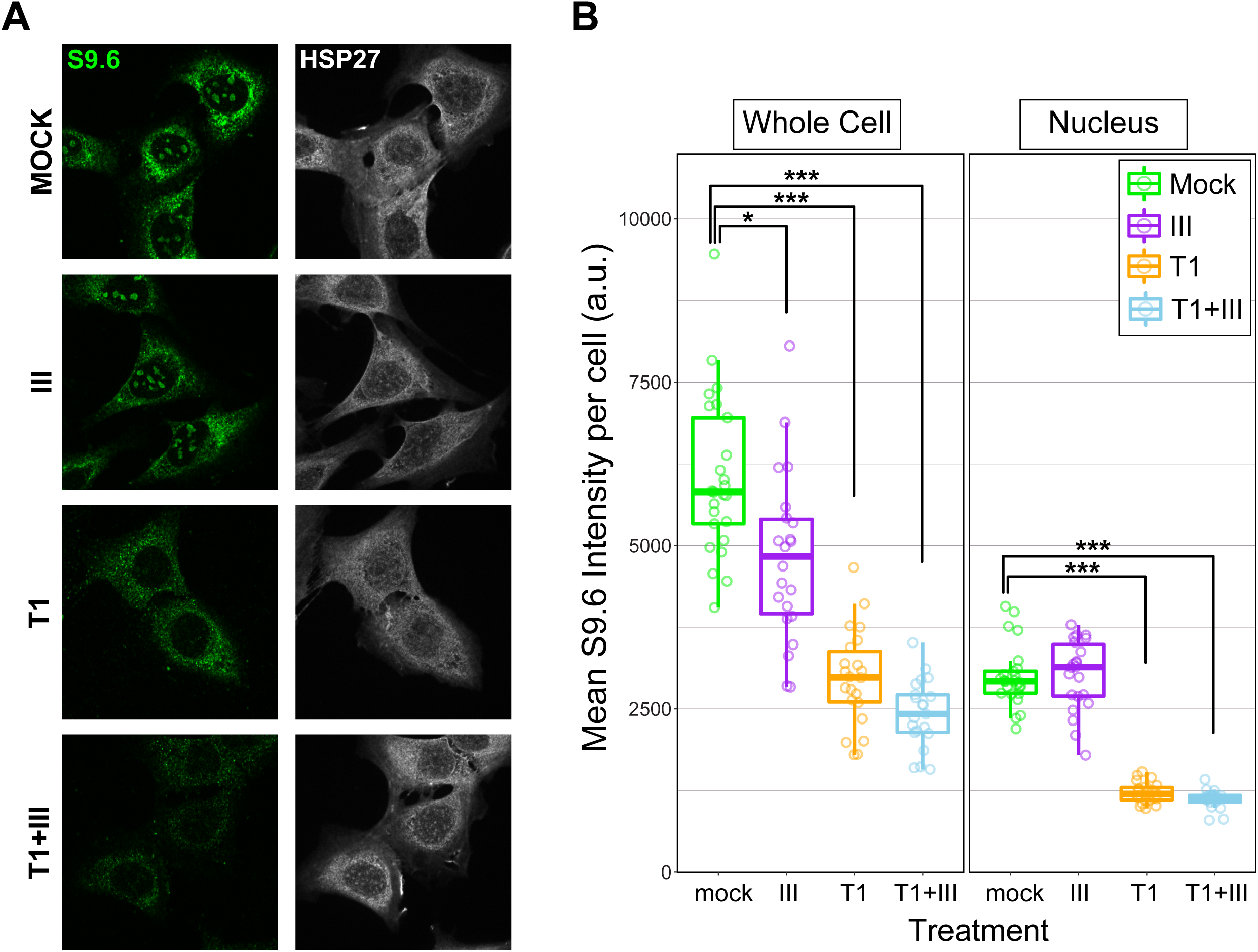
**A.** Representative images of single planes of formaldehyde-fixed U2OS cells that were mock-treated or pre-treated RNase III, RNase T1, or a combination of both enzymes post-fixation for 1 hour at room temperature and stained with S9.6 (green) and anti-HSP27 (white). **B.** Quantification of whole cell and nuclear mean S9.6 intensities for individual cells that were mock- or enzyme-treated.

**FIGURE S4:**
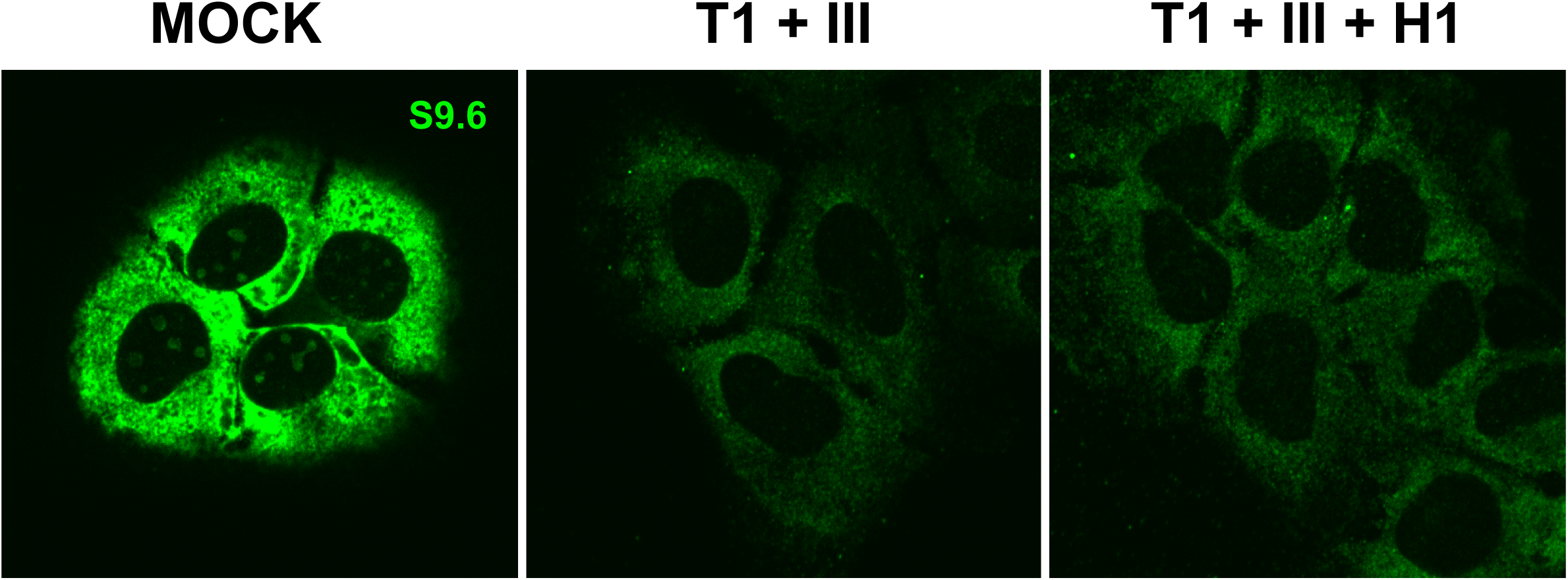
Images of single planes of methanol-fixed U2OS cells labeled with S9.6 that were mock-treated or pre-treated with a combination of RNase T1 and III and a combination of RNase T1, III, and H1.

**FIGURE S5:**
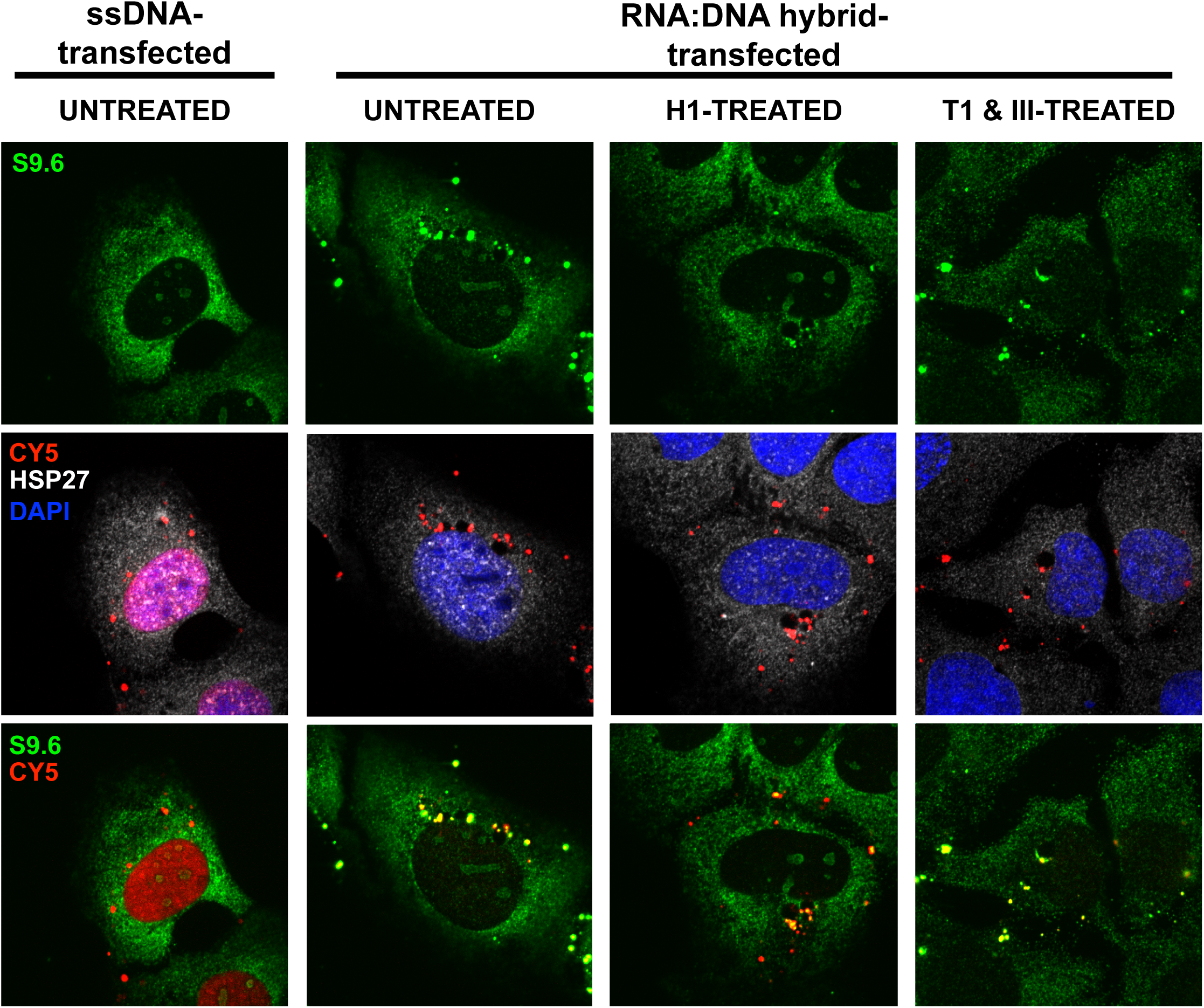
Images of single planes of U2OS cells transfected with 5’-Cy5-labeled ssDNA and RNA:DNA hybrids (red) and then fixed and immunolabeled with S9.6 (green) and anti-HSP27 (white). RNA:DNA hybrid transfected cells were mock-treated and pre-treated with RNase H1 and a combination of RNase T1 and III.

## ACKNOWLEDGEMENTS

Work in the Chedin lab is funded by a grant from the National Institutes of Health (R01 GM 120607). J.S. was funded in part through the NIH T32 training program in Molecular and Cellular Biology (T3 GM 007377). We thank Drs. Frank McNally, Jodi Nunnari and Samantha Lewis for assistance with the work and useful suggestions.

